# Ensemble gene function prediction database reveals genes important for complex I formation in Arabidopsis thaliana

**DOI:** 10.1101/181396

**Authors:** Bjoern Oest Hansen, Etienne H. Meyer, Camilla Ferrari, Neha Vaid, Sara Movahedi, Klaas Vandepoele, Zoran Nikoloski, Marek Mutwil

**Affiliations:** Max Planck Institute of Molecular Plant Physiology, Am Muehlenberg 1, 14476 Potsdam, Germany; Institut für Medizinische Informatik, Universitätsmedizin Göttingen, Robert-Koch-Str. 40 37075 Göttingen, Germany; VIB Center for Plant Systems Biology, Department of Plant Biotechnology and Bioinformatics Ghent University, Technologiepark 927, B-9052 Gent, Belgium; Rijk Zwaan Breeding B.V., Burgemeester Crezéelaan 40, PO Box 40, 2678 ZG De Lier, The Netherlands; Bioinformatics Group, Institute of Biochemistry and Biology, University of Potsdam, Karl-Liebknecht-Str. 24-25, 14476 Potsdam-Golm, Germany; School of Biological Sciences, Nanyang Technological University, 60 Nanyang Drive, Singapore 637551, Singapore

**Keywords:** gene function prediction, ensemble prediction, co-function network, complex I, *Arabidopsis thaliana*

## Abstract

- Recent advances in gene function prediction rely on ensemble approaches that integrate the results from multiple inference methods to produce superior predictions. Yet, these developments remain largely unexplored in plants.
- We have explored and compared two methods to integrate ten gene co-function networks for *Arabidopsis thaliana* and demonstrate how integration of these networks produces more accurate gene function predictions for a larger fraction of genes with unknown function.
- These predictions were used to identify genes involved in mitochondrial complex I formation, and for five of them we confirmed the predictions experimentally. The ensemble predictions are provided as a user-friendly online database, EnsembleNet, available at http://www.gene2function.de/ensemblenet.html.
- The methods presented here demonstrate that ensemble gene function prediction is a powerful method to boost prediction performance, while the EnsembleNet database provides cutting-edge community tool to guide experimentalists.

## Introduction

Extracting useful knowledge from genomes is dependent on our ability to correctly assign biological functions to gene products. However, since proteins may have more than one function that are carried out under different condition (e.g. developmental stage, environment), experimental characterization of protein function is resource-intensive. Therefore, the challenge posed by the sheer number of uncharacterized gene products has rendered gene function prediction as one of the most pressing bioinformatics research efforts (Radivojac *et al.*, 2013; Jiang *et al.*, 2016, reviewed in Rhee & Mutwil, 2014). Apart from providing an overview of the functional repertoire of an organism, gene function predictions in plants are invaluable for experimentalist, as they provide the basis for a rapid reverse genetic approach to identify relevant genes. For example, several studies used *in silico* predictions to identify genes involved in plant viability (Mutwil *et al.*, 2010), seed germination (Bassel *et al.*, 2011), shade avoidance (Jiménez-Gómez *et al.*, 2010), cyclic electron flow (Takabayashi *et al.*, 2009), cell division (Takahashi *et al.*, 2008), drought sensitivity and lateral root development (Lee *et al.*, 2010).

Current protein function prediction methods largely rely on the Gene Ontology (Ashburner *et al.*, 2001) to communicate and describe functional annotation, and to describe relationships between those annotations. The Gene Ontology (GO) uses three functional domains to specify gene function: Cellular Component (CC, subcellular component/protein complex where the gene product acts), Molecular Function (MF, biochemical activity of the gene product), and Biological Process (BP, purpose of the gene product). For example, *Atlg72370*, a 40S ribosomal protein SA (Garcia-Hernandez *et al.*, 1994), is an “mRNA binding” protein (GO:0003729, MF), which is part of “cytosolic small ribosomal subunit” (GO:0022627, CC), and is involved in “mature ribosome assembly” (GO:0042256, BP, www.arabidopsis.org). Importantly, GO also provides evidence codes useful in specifying if an annotation is based on an experiment (e.g. IEP: inferred from expression pattern) or a prediction (e.g. ISA: inferred from sequence alignment), which is needed to construct a “gold standard” dataset of experimentally verified genes, used in turn for propagating function from characterized to uncharacterized genes (Radivojac *et al.*, 2013; Jiang *et al.*, 2016, reviewed in Rhee & Mutwil, 2014).

Gene function prediction, regardless of the data used, requires three components: (i) a dataset that specifies features of the analyzed genes, (ii) a method to identify functionally related genes in the given dataset and (iii) a method to transfer functional annotation from the gold standard to uncharacterized genes (Radivojac *et al.*, 2013; Jiang *et al.*, 2016, reviewed in Rhee & Mutwil, 2014). For example, sequence homology-based function predictions (e.g. InterProScan, Quevillon *et al.*, 2005) use (i) a database of functionally annotated protein sequences (the gold standard), to (ii) find functionally related proteins that are identified via BLAST, with the aim to (iii) transfer functional information from a similar gold standard protein to an uncharacterized protein. The gene function prediction methods can be divided into three “generations”: (i) single, (ii) integrative and (iii) ensemble (reviewed in Proost & Mutwil, 2016). However, regardless of the prediction method, all approaches are based on the guilt-by-association (GBA) principle, whereby genes with similar features are likely to have same function (Pavlidis & Gillis, 2012, reviewed in Rhee & Mutwil, 2014).

First generation methods use a single type of data to identify functionally related genes (reviewed in Rhee & Mutwil, 2014). Genomic GBA approaches use sequence similarity (e.g. InterProScan, Quevillon *et al.*, 2005), genomic co-occurrence (i.e. gene families that are co-gained or co-lost across species have same function, Pellegrini *et al.*, 1999), genomic neighbor (i.e. proximity on chromosome implies same function, Lee & Sonnhammer, 2003), gene fusion (i.e. two separate gene products that are found as one protein in a species, are interacting, Nakamura *et al.*, 2007 and domain co-occurrence (i.e. protein domains that frequently co-occur on proteins are functionally related, Wang *et al.*, 2011 to identify functionally related genes. Transcriptomic GBA approaches utilize co-response to perturbation and co-expression to predict gene function. For instance, transcripts that are up-regulated upon cold stress may be involved in cold adaptation (Lee, 2005) and functionally related genes tend to show similar expression profiles across treatments (Wu *et al.*, 2002). Similarly, protein-protein interaction GBA states that interaction of two proteins generally implicates that they are functionally related (e.g. subunits of a ribosome, Vazquez *et al.*, 2003).

Second generation methods integrate different data types, thereby addressing important shortcomings of the first generation methods (Heyndrickx & Vandepoele, 2012a, reviewed in Rhee & Mutwil, 2014). These shortcomings include limited coverage (e.g. the Arabidopsis ATH1 microarray captures only 60% of protein coding transcripts, Mutwil *et al.*, 2011, or <10% of the predicted Arabidopsis interactome has been elucidated, Arabidopsis Interactome Mapping Consortium, 2011), technical artifacts (e.g. bimolecular fluorescence complementation (BiFC) interaction studies produce false positive results due to protein over-expression, Kudla & Bock, 2016) and technical limitation (e.g. yeast-two-hybrid (Y2H) only works on proteins that can be imported into the nucleus). Integration of different datasets can result in increased coverage (e.g. combining BiFC and Y2H can increase the size of inferred interactome) and confidence (e.g. protein interactions reported by both BiFC and Y2H are less likely to be false positives). Indeed, through Bayesian integration of protein-protein interactions, literature text mining, co-expression, genomic neighbor, gene function and domain co-occurrence, Lee and colleagues constructed a co-function network with prediction accuracy higher than any of the single methods (Lee *et al.*, 2010). Furthermore, since different subset of biological information is captured by each method, integrating the diverse biological data can result in a more complete functional description (Heyndrickx & Vandepoele, 2012b).

Third generation ensemble (also called community) methods produce a prediction by integrating the outputs of multiple first and second generation methods. The concept is similar to the “Ask the audience” lifeline (91% success rate) in the Who Wants to Be a Millionaire television quiz, which performs better than “phone a friend” (65% success rate) lifeline (“The Wisdom of Crowds”, James Surowiecki, 2004). Indeed, integration of 29 gene regulatory network inference methods in yeast and *E. coli* generated an ensemble prediction that outperformed all of the individual methods (Marbach *et al.*, 2012). The ensemble from as few as three input networks emphasized the advantages and diminished the limitations of each of the 29 methods.

In plants, ensemble approaches are largely unexplored. SUBA3 provides ensemble sub-cellular predictions based on 22 prediction programs (Tanz *et al.*, 2013). Furthermore, by integrating four different methods, a recent study predicted an ensemble gene regulatory network in Arabidopsis (Vermeirssen *et al.*, 2014). To our knowledge, ensemble gene function predictions are not available for *Arabidopsis thaliana* or other plants, which we set to remedy in this study.

## Materials and Methods

### Retrieval and normalization of gene co-function networks

The most recent versions of networks available for *Arabidopsis thaliana* on 21.11.2015 were downloaded for AraNet2 (http://www.inetbio.org/aranet/downloadnetwork.php), STRING (https://string-db.org/cgi/download.pl), GeneMANIA (http://genemania.org/data/), ATTED-II (http://atted.jp/download.shtml), PlaNet (http://aranet.mpimp-golm.mpg.de/download.html), Conserved Co-expression Arabidopsis Functional Modules (CC-AFM), PODC (http://plantomics.mind.meiji.ac.jp/podc/bulkDownload.html), TAIR-PPI (ftp://ftp.arabidopsis.org/home/tair/Proteins/Protein interaction data/) and BIOGRID (https://thebiogrid.org/download.php). CC-AFM refers to Arabidopsis co-expression edges (top 300 per gene based on Pearson Correlation Coefficient) filtered with conserved coexpression information from six other plants, including *Glycine max, Medicago truncatula, Oryza sativa, Populus trichocarpa, Vitis vinifera* and *Zea mays* from (Heyndrickx & Vandepoele, 2012b).

Furthermore, normalized expression data derived from RNA-seq experiments from (Giorgi *et al.*, 2013) was used to construct Highest Reciprocal Rank (HRR) co-expression networks, as done in PlaNet (HRR cut-off=30, Mutwil *et al.*, 2010).

Edges of each of the networks were scaled to range between 0 and 1, where 1 indicates the strongest association between a gene pair, while 0 indicates the weakest association.

### Retrieval of gold standard - experimentally verified functional annotation

Experimentally verified Gene Ontology annotations were downloaded from TAIR (www.arabidopsis.org), as ATH_GO_GOSLIM.txt file on 21.11.2015 (used to construct and benchmark EnsembleNet) and 10.08.2017 (used to perform time-stamp analysis). A gene was considered experimentally annotated if it was associated with GO terms having EXP, IDA, IPI, IMP, IGI, IEP or TAS evidence codes. To avoid evaluating general terms, (e.g. “cellular_component”, “binding”, ”biological regulation”), terms that were assigned to more than 5% of the 14,031 experimentally annotated genes were removed. This removed 105 out of 7,352 terms. For each gene, all terms assigned to a gene were recursively propagated towards the root of the terms (excluding the terms present in more than 5% of the experimentally annotated genes), regardless of the type of relationship between terms.

### Calculating F-measure

The main evaluation metric in this study was F-measure, as defined by Critical Assessment of protein Function Annotation (CAFA) experiment (Radivojac *et al.*, 2013; Jiang *et al.*, 2016).

For a given target gene *i* and network edge threshold *t* ∈ [0,1], the precision and recall were calculated as:

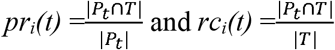

where *|P_t_|* is the number of predicted terms at edge threshold *t*, |T| is the number of experimentally determined terms, and |P_t_∩T| is the number of terms appearing in both P_t_ and T.

Performance of a network at a given edge threshold is obtained by averaging precision and recall across genes. Precision at edge threshold *t* is calculated as:

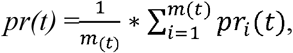

where *m(t)* is the number of genes for which at least one prediction was made, above edge threshold t. Precision is calculated only for the genes for which a prediction could be made. A prediction cannot be made for genes that are not connected to genes with any experimentally determined annotations at a given *t*.

Conversely, recall is calculated over all *n* genes in a target set, as:

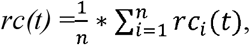

regardless of whether a prediction can be made at a given threshold *t*.

The F-measure (a harmonic mean between precision and recall) provides a number useful to compare performances of the networks at a given threshold. An edge threshold where a given network performs highest in terms of the F-measure, can be identified by obtaining the maximum F-measure (*F_max_*) over all thresholds, given by:

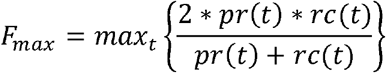

### Associating genes to Gene Ontology terms

To predict which genes are associated with a given GO term (query term), we first removed GO terms assigned to more genes that the query term from the gold standard. The reason for removing these terms is that more frequent terms (e.g. “binding”, “enzymatic activity”) are by chance more likely to be more abundant in network neighborhoods than less frequent terms. Next, for each gene in the used networks (i.e. GeneMANIA and AraNet v2 for MF domain), NCE was used to predict genes with the query term.

### Calculating the empirical p-value to evaluate gene - Gene Ontology term associations

To estimate how accurately genes assigned to a given Gene Ontology term can be predicted by NCE, we first retrieved genes experimentally assigned to the term of interest. Then, for each of these genes, we counted the number of networks for which the neighbor counting made a correct prediction (Figure S2). This number, pred_observed_, is indicative of the extent to which a term can be accurately predicted in a network setting. See Figure S5A and S5B for toy examples where a term can be predicted well (predobserved = 9, out of 9 predictions) and poorly (pred_observed_ = 2, out of 9 predictions), respectively.

Next, for each gene that is experimentally assigned to the term, we sampled a random prediction from the pool of all predicted gene-term associations, and counted the number of networks which assigned a term to a gene (pred_sampled_). The empirical p-value is defined as:

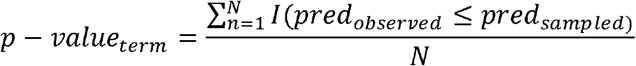

Where predobserved is the observed number of correct predictions, predsampled is the number of correct calls from sampled predictions, N is the number of permutations, and I is an indicator function. To account for multiple hypothesis testing, we applied False Discovery Rate (FDR) correction to the p-values (Benjamini & Hochberg, 1995). The predicted gene-GO term associations and p-values are available as Supplemental Data 1.

### Calculating STATIS compromise network

To combine the 10 symmetric (i.e., undirected) weighted networks, we applied an approach similar to STATIS (Abdi *et al.*, 2012). We first determined the Rv coefficient for each pair of the scaled networks (represented as scaled adjacency matrices), gathered in the cosine matrix C. The entries in the first eigenvector of C, after rescaling to unit norm, were used as weights to determine the contribution of each scaled adjacency matrix to the STATIS network. We note that the larger the value for the weight, the bigger the contribution of the respective data set to the adjacency matrix of the STATIS network. The STATIS network was obtained as the weighted sum of the scaled adjacency matrices. The resulting adjacency matrix *S* was also scaled by dividing with its maximum value. The STATIS network is available for download at http://aranet.mpimp-golm.mpg.de/download.html.

### Identifying functionally related genes

To estimate which genes are functionally related, we first retrieved the predicted terms for each gene, and calculated the Jaccard Index between all possible gene pairs, defined as: J(X,Y) = |X∩Y| / |X∪Y|, where X and Y are predicted GO terms of gene x and y. To estimate which gene pairs are functionally related, we calculated the Jaccard Index between 10.000 random gene pairs and found the J threshold (0.178) below which 99% of the random pairs were found (Figure S6). Thus, gene pairs with J>0.178 were deemed to be functionally related. The resulting GOJI network is available as Supplemental Data 2.

### Plant material, plant growth and genotyping

Seeds were surface sterilized with 70 % (v/v) ethanol containing 0.5 % (v/v) Triton X100, sown on synthetic media (0.5× MS, 1 % sucrose, 0.7 % agar) and incubated under long day photoperiod: 16 h light (150 μE, 22 °C)/8 h dark (20 °C). After 2 weeks, seedlings were transferred to soil and grown under long day photoperiod. Plant DNA was isolated as described in (Weigel & Glazebrook, 2009), and genotyping PCR was performed as in (O’Malley *et al.*, 2015). The genotyping primers for detecting the wild type and mutant alleles are given in Table S8.

### Mitochondria isolation, Blue-Native gel electrophoresis and quantification of complex I

The aerial organs of 6-week-old plants were harvested and mitochondria were isolated according to (Kühn *et al.*, 2015). For Blue-Native gel electrophoresis, total mitochondria proteins (50 μg) were solubilized with dodecylmaltoside (1% [w/v] final) in ACA buffer (750 mM amino caproic acid, 0.5 mM EDTA, and 50 mM Tris-HCl, pH 7.0) and incubated 20 min at 4°C. After centrifugation for 10 min at 20000g, the supernatant was recovered and supplemented with Coomassie Blue G250 (0.2 % [w/v] final). The samples were loaded onto a 4.5% to 16% (w/v) polyacrylamide gel (ratio acrylamide/bisacrylamide: 29/1, gel buffer: 0.25 M aminocaproic acid, 25 mM Bis-Tris-HCl pH 7.0). The migration of the samples was performed at 4°C using cathode buffer (50 mM Tricine, 15 mM Bis-Tris-HCl pH 7.0, 0.02% [w/v] Coomassie Blue G250) and anode buffer (50 mM Bis-Tris-HCl pH 7.0) in two steps, first 100 V for 45 min then 15 mA (limiting voltage: 500 V) for 10 h. After migration the gel was stained using colloidal blue (3% [v/v] orthophosphoric acid, 6% [w/v] ammonium sulphate, 0.1% [w/v] Coomassie Blue G250) and destained with ultrapure water. The gel was scanned using a flatbed scanner. The gel analyzer tool from ImageJ was used to quantify the bands corresponding to complexes I, III and V.

## RESULTS

### Benchmarking of ten gene co-function networks

In this study, we obtained and standardized ten gene co-function networks (see Methods). AraNet v2, GeneMANIA and STRING are second generation methods integrating multiple data sources (Table 1). The remaining seven networks are obtained from first-generation methods based on co-expression (ATTED-II, CC-AFM, PlaNet-microarrays, PlaNet-RNAseq, PODC) or protein-protein interactions (BIOGRID, TAIR-PPI). To determine and evaluate the predictions, we have obtained the gold standard Gene Ontology annotations, based on experimental evidence for 14,032 *Arabidopsis thaliana* genes (www.arabidopsis.org, Lamesch *et al.*, 2012). Analysis of the gold standard revealed that 31% of the 28,392 Arabidopsis genes in BP domain have been experimentally characterized (Figure S1A), while less than 12% of genes have all three GO domains experimentally characterized (Figure S1B).

**Table 1.**
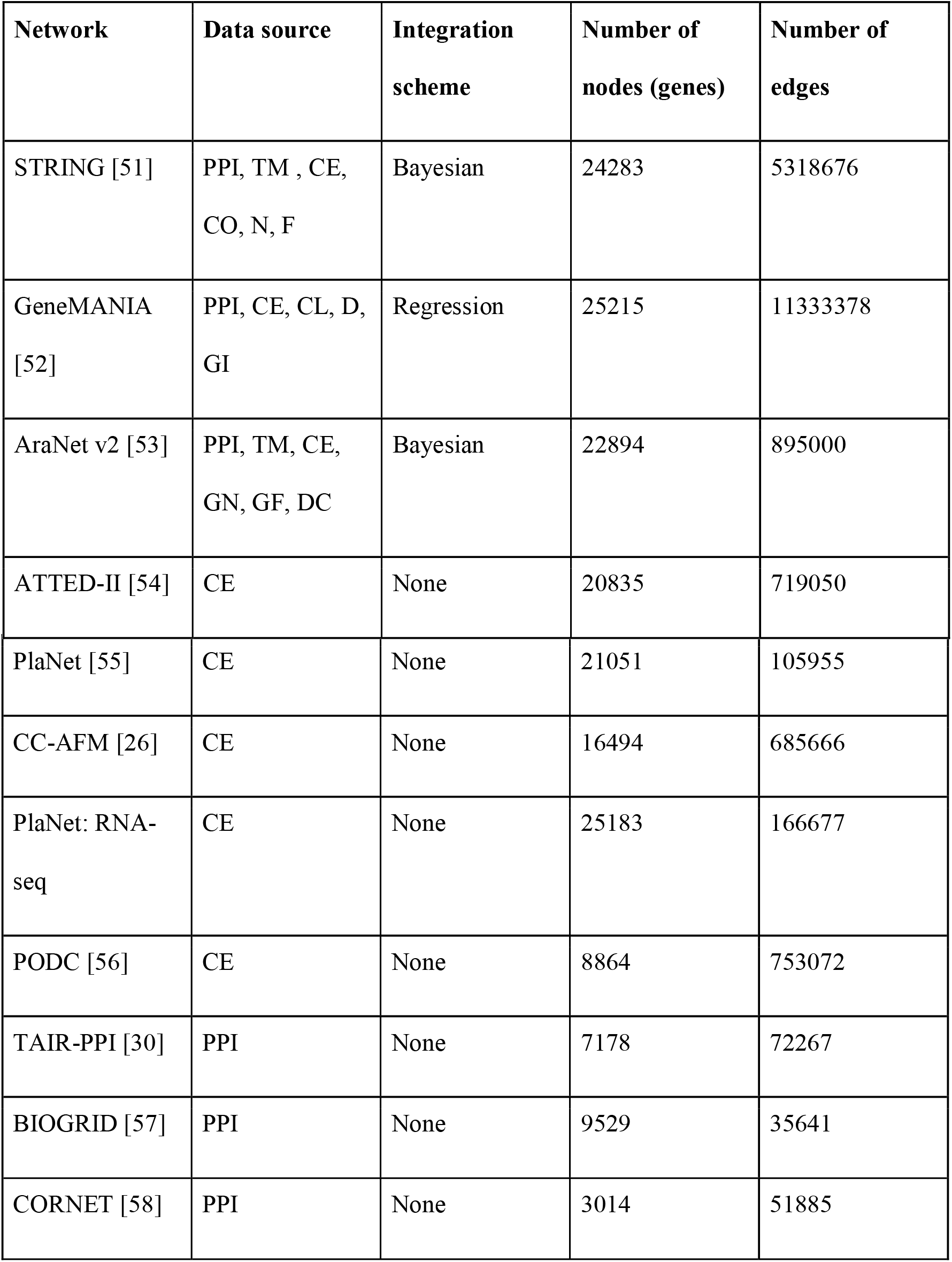
Used co-function networks. Type of data and integration scheme. PPI (protein-protein interaction), TM (text mining), CE (co-expression), CO (genomic co-occurrence), GN (genomic neighbor), GF (gene fusion), CL (co-localization), D (shared protein domains), GI (genetic interaction), DC (domain cooccurrence).

To predict gene function, we used the Neighbor Counting (NC) approach as the simplest method for network-based function prediction (Sharan *et al.*, 2007). NC predicts function of a gene by identifying the most frequent function(s) found in the network neighborhood of the gene (Figure 1A). Furthermore, since all networks (apart from PPI networks) contain edge weights that can be scaled to range from 0 (weakest association between genes) to 1 (strongest association), a network can perform differently at a specific edge cut-off. The performance of a network at a given edge cut-off is specified by the performance of the subnetwork obtained upon removal of all edges with weight smaller than the specified cut-off. Therefore, too low an edge cut-off could connect functionally unrelated genes, resulting in an incorrect prediction (Figure 1A, first network), or partially correct prediction (Figure 1A, second network). Conversely, too high a cut-off could disconnect functionally related genes and thus hinder any prediction (Figure 1A, fourth network).

**Figure 1.**
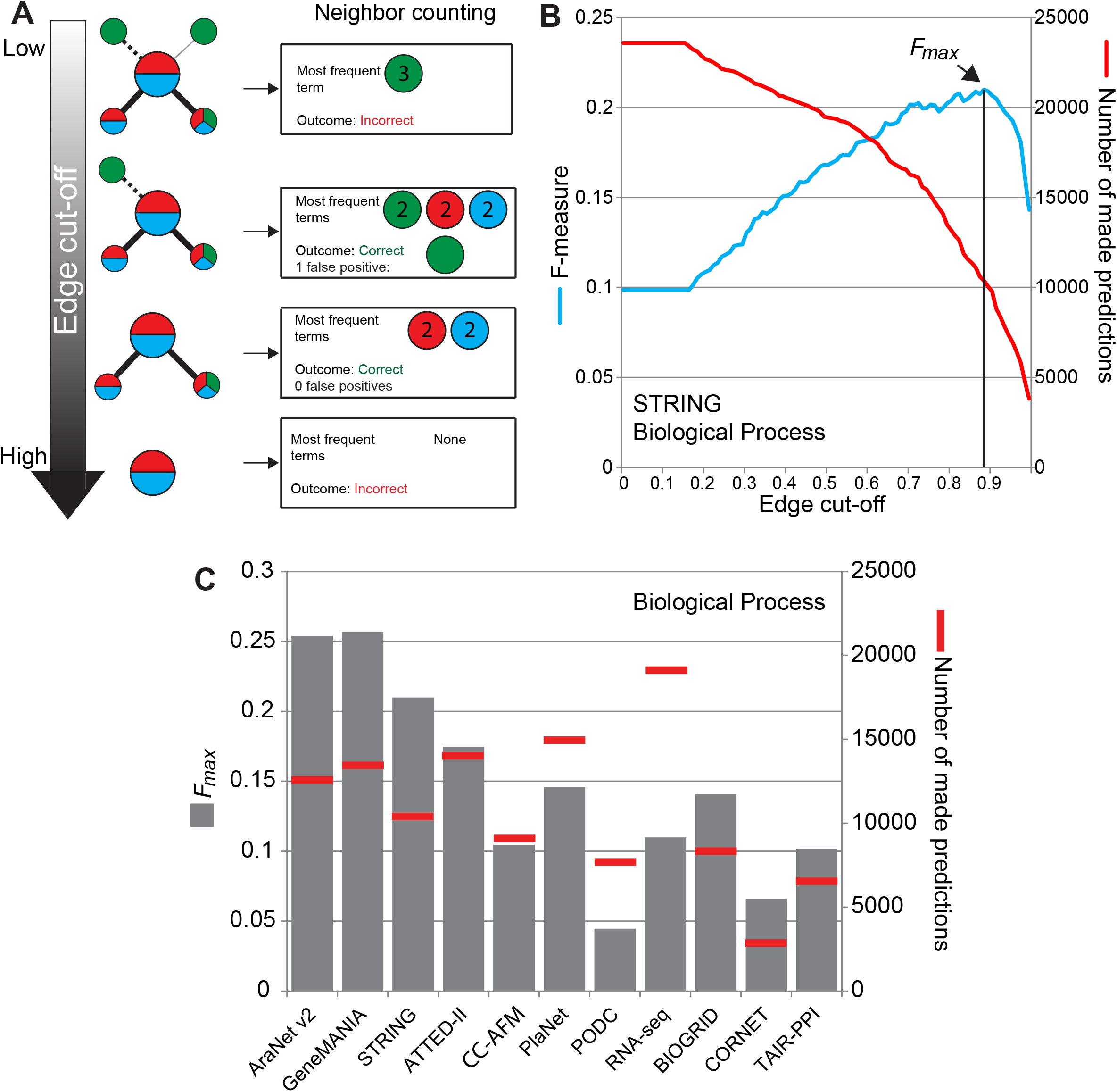
Evaluation of the ten gene co-function networks. A) Illustration of the Neighbor Counting method and influence of the edge cutoff on predictions, exemplified by four toy examples. The toy examples include the query gene (large node) connected to other genes in the co-function network (smaller nodes). The edges connecting genes have weights ranging from weak (solid gray edge) to strong (solid black edge). Gene functions assigned to a gene are depicted as red, blue and green colors. Neighbor Counting method assigns function to the query gene (large node) that occur most frequently in its neighbors. The accuracy of the prediction can be estimated by comparing the known functions of the query with the predicted function. B) Influence of the edge cutoff on the F-measure (blue line) and made predictions (red line) for STRING BP predictions. The edge cutoff where F-measure is highest (*F_max_*) is indicated. C) Performance of the ten co-function network expressed as *F_max_* value (gray barcharts, left y-axis) and the number of made predictions (red bars, right y-axis).

To provide a fair evaluation and comparison of the networks, we first set to find an edge threshold where a given network produces the highest number of correct predictions. To this end, we used the F-measure which represents the harmonic mean between precision (i.e. average fraction of correctly predicted terms) and recall (i.e. average fraction of correctly predicted terms over all relevant terms, Radivojac *et al.*, 2013), as a measure of performance for prediction methods in Critical Assessment of Functional Annotation (Radivojac *et al.*, 2013). We applied a leave-one-out cross validation, where the function of an experimentally characterized gene is compared to the Neighbor Counting-predicted function of the gene, treating its function as unknown. The F-measure can range from 0 (worst value) to 1 (best value) and provides a number that can be used to compare the performance of the networks (Table S1A).

We illustrate this type of analysis with the STRING network, for which we evaluated the performance for the BP domain at a given edge cut-off. Not surprisingly, as a consequence of genes becoming disconnected in the network, the number of made predictions dropped as the edge cut-off increased (Figure 1B, red line, right y-axis). The F-measure peaked at 0.88 edge cut-off, identifying the *F_max_* value where STRING produces highest prediction accuracy for BP domain (Figure 1B, blue line, left y-axis). Interestingly, *F_max_* for the BP, CC and MF domain were found at 0.88, 0.6 and 0.65, respectively, suggesting that one edge cut-off may not be suitable to produce most accurate predictions for the three domains (Table S1B).

To compare the 10 co-function networks, we plotted the *F_max_* values and the number of made predictions (Figure 1C). As expected from networks integrating multiple data sources, AraNet v2, GeneMANIA and STRING showed higher *F_max_* values than coexpression and PPI-based methods (Figure 1C, gray bars, left y-axis), but the two coexpression networks with highest *F_max_* (ATTED-II and PlaNet) showed higher number of made predictions (Figure 1C, red bars, right y-axis).

### Generation and benchmarking of ensemble gene function predictions

To construct community predictions from the 10 co-function networks, we explored two different approaches to integrate these networks: the first approach, termed Neighbor Counting Ensemble (NCE) expands the NC rule to multiple networks. More precisely, NCE (i) uses Neighbor Counting to predict function of a gene in each of the input networks and (ii) identifies the most frequent term found by NC in the previous step (ensemble majority vote, Figure 2A). To investigate which combination of the co-function networks produces *F_max_* for NCE, we tested all combinations of 2 to 19 networks at their *F_max_* threshold, resulting in 1,023 tested combinations for each GO domain (Table S2).

**Figure 2.**
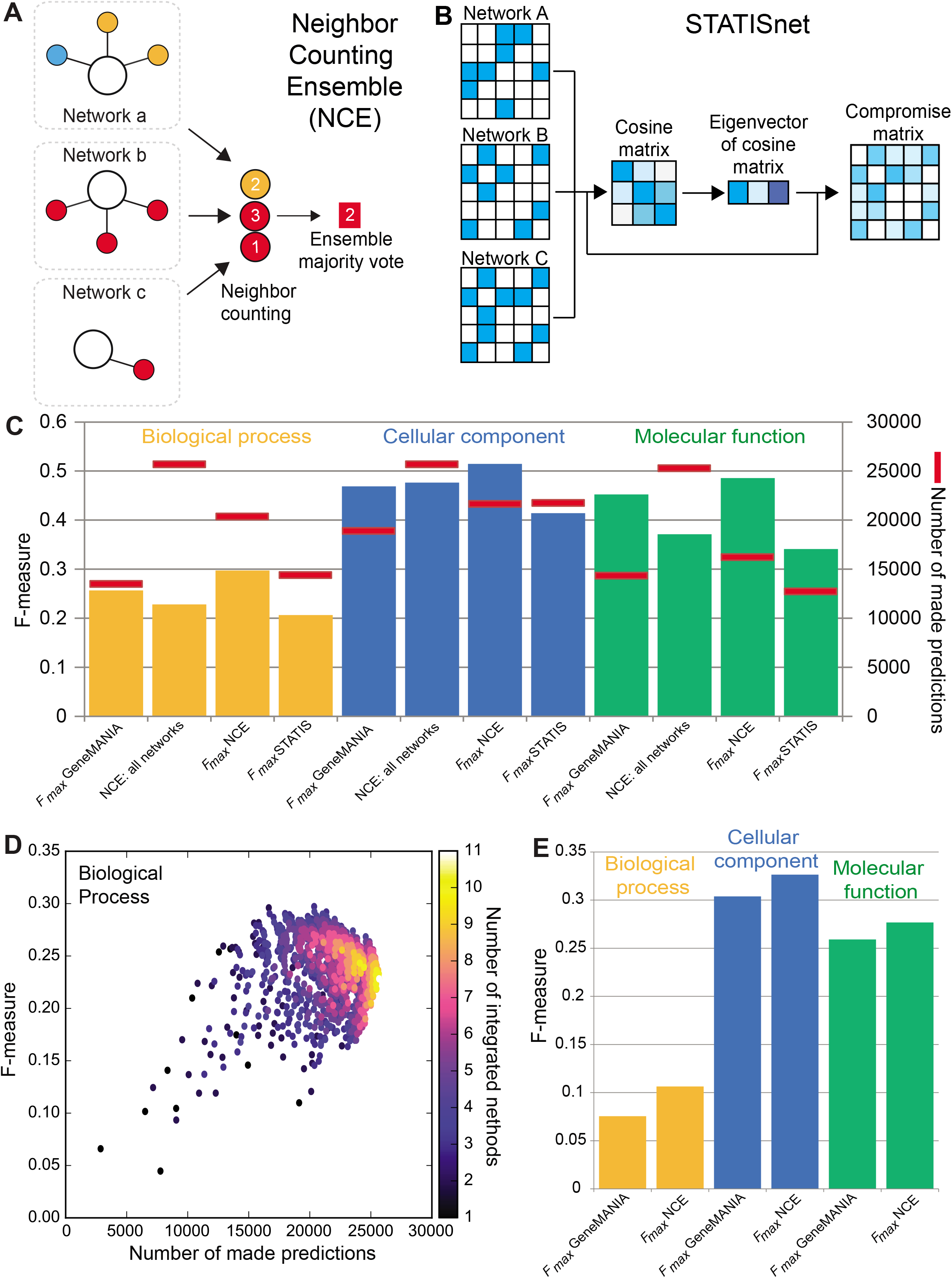
Performance of ensemble gene function prediction methods. A) Outline of the Neighbor Counting Ensemble (NCE) approach, exemplified by a toy example. Function for the query gene (large white node) is estimated by neighbor counting, predicting orange, red and red function from networks a, b and c, respectively. In the next step, similarly to the Neighbor Counting, the ensemble prediction is obtained by selecting the most frequent function from the previous step, assigning red function to the query gene. B) Outline of the STATIS method. C) Performance summary of the methods for BP (red), CC (blue) and MF (green). The left y-axis (bars) indicates the F-measure, while the right y-axis (black line) indicates the number of made predictions. D) Analysis of the 10 gene co-function networks, resulting in 1,023 tested combinations. The x-axis indicates the number of made predictions, while the y-axis shows the F-measure. Point color indicates the number of integrated networks, ranging from 1 network (dark blue) to 10 (bright yellow). E) Time-stamp comparison of the Ensemble method and GeneMANIA predictions. The y-axis indicates the F-measure.

The second approach, based on STATIS (Abdi *et al.*, 2012), termed STATISnet, determines the similarity between each pair of the scaled network adjacency matrices, by calculating Rv coefficient of the adjacency matrices. The principal eigenvector is then used to build a weighted sum of the scaled adjacency matrices, which yields the adjacency matrix of STATISnet (Figure 2B). The *F_max_* for STATISnet was then calculated as for the 10 input networks (Table S1).

The analysis revealed that NCE produced highest *F_max_* and number of made predictions (prediction coverage) than the best single method (GeneMANIA), and STATISnet for all three GO domains (Figure 2C, Table S2). While *F_max_* of STATISnet was lower than *F_max_* of GeneMANIA, the prediction coverage of the former was modestly higher for BP and CC domains (Figure 2C). Interestingly, NCE based on all 10 networks produced a lower value for the F-measure than NCE based on GeneMANIA, AraNet v2 and ATTED-II, indicating that the combination of selected networks may have a negative effect on the prediction performance (Table S2). Furthermore, while *F_max_* NCE for BP and CC domains used GeneMANIA, AraNet v2 and ATTED-II, NCE for MF utilized GeneMANIA and AraNet v2, suggesting that different types of information may increase prediction accuracy of different GO domains.

To further gain insight into how network selection can influence NCE predictions, we plotted the F-measures of the 1,023 tested combinations (y-axis, Figure 2D) versus the number of made predictions for each combination (x-axis). The plot revealed that while integrating all 10 co-function networks increases the number of made predictions (Figure 2D, bright yellow color), the combinations that produce high F-measures included fewer than 6 networks (Figure 2D, blue color), which was observed for the three GO domains (Figure S2, Table S2).

In addition to the leave-one-out cross validation to benchmark the networks, we also performed a timestamp analysis (Radivojac *et al.*, 2013; Jiang *et al.*, 2016), where predictions based on gold standard from November 2015 were compared to functional information available on June 2017. Comparison of these two gold standards revealed that within 20 months, 488 previously unannotated genes gained functional information, while 2,588 genes gained additional GO terms. We then calculated the F-measure of our predictions based on the new functional information. Similarly to the previous comparison, the timestamp analysis revealed that NCE had higher F-measure than the best single performing method (GeneMANIA) for all three GO domains (Figure 2E, Table S3).

### Influence of the gold standard completeness on the prediction accuracy and coverage

Since the gold standard is used to propagate functional information to uncharacterized genes in gene function prediction studies, the size of the gold standard should strongly influence the quality and number of predictions.

We first investigated how the amount of available functional information influences the accuracy of the predictions. To this end, we incrementally removed 10% randomly selected genes from the gold standard of 14,032 genes, and calculated the F-measure for the remaining genes with leave-one-out analysis. Not surprisingly, when only 10% (1,403) of gold standard genes retained functional information, the mean value for the F-measure over 100 repetitions decreased for NCE and AraNet v2, GeneMANIA and STRING (Figure 3A). However, NCE and GeneMANIA, and to a lower degree AraNet v2 and STRING, approached their highest F-measures at gold standard containing ~50% of genes (Figure 3A). Since NC uses network neighbors to predict gene function, the number of made predictions showed strong dependency on the size of the gold standard (Figure 3B). This was also observed for the CC and MF domains (Figure S3). Taken together, the completeness of the gold standard has a strong effect on the number and the accuracy of made predictions.

**Figure 3.**
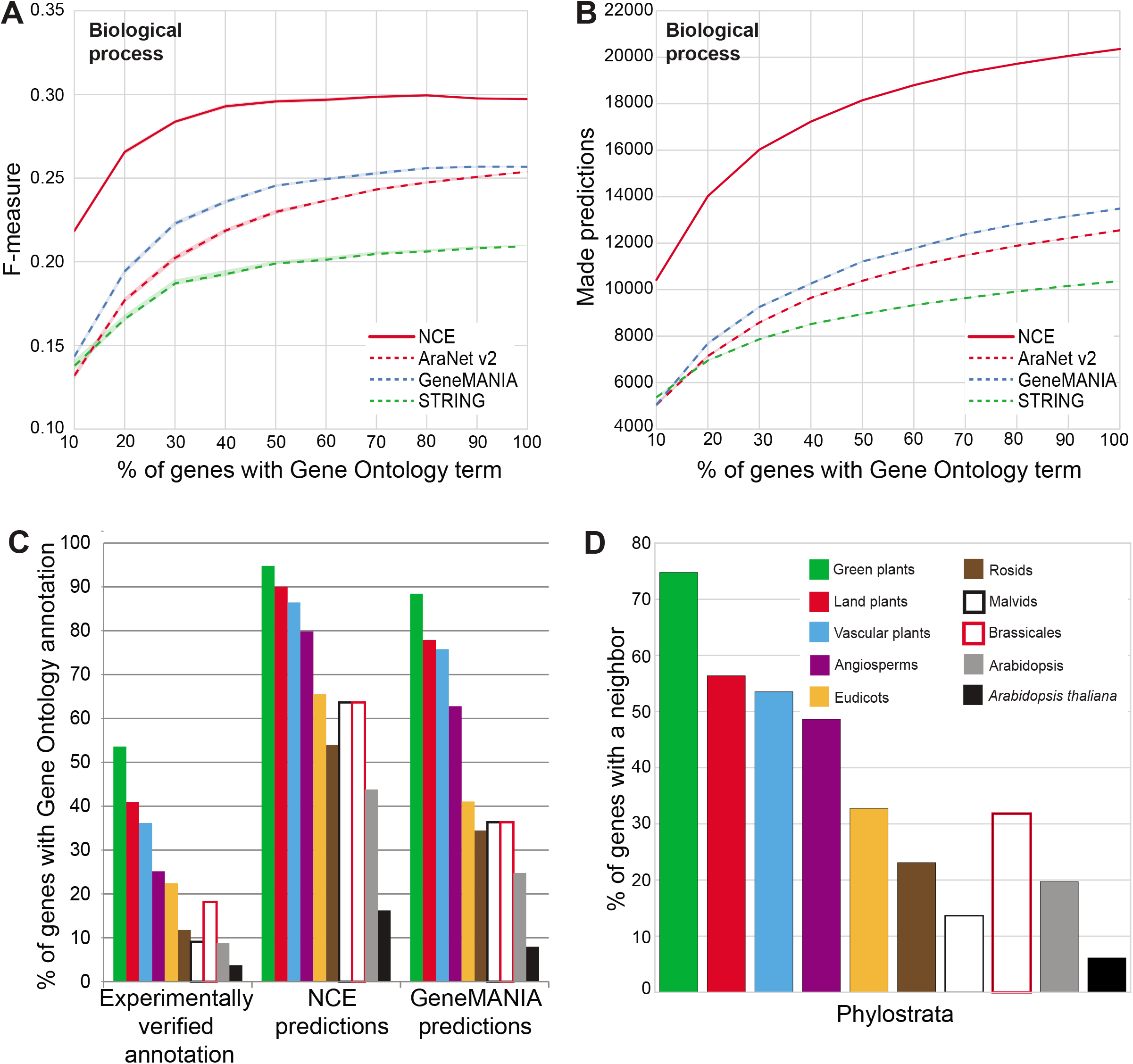
Influence of available functional information on prediction performance. A) Relationship between the F-measure (y-axis) and % of genes with available GO terms (x-axis) in the three best performing co-function networks (dashed lines) and NCE (solid red line). The lines represent bootstrapped 68% confidence interval plots of 100 permutations. B) Relationship between the number of made predictions (y-axis) and % of genes with available GO terms (x-axis). C) Percentage (y-axis) of genes from the different phylostrata (given by color legend in Figure 3D), for experimentally characterized genes, Ensemble predictions and GeneMANIA predictions. D) Relationship between the percentage of genes with a neighbor in a network (y-axis) and the phylostrata of the genes (x-axis). The example is given for GeneMANIA, BP.

### Negative correlation between gene age and gene function predictability

Our previous analyses revealed that younger genes tend to be less functionally characterized than older genes, as the gold standard is composed mostly of older gene families (Ruprecht *et al.*, 2017). While the cause for this phenomenon can only be speculated (e.g. younger genes have weaker, easy to miss mutant phenotypes), the amount of functional information almost perfectly reflects the order of appearance of the gene families. We ordered the gene families from oldest (i.e. Green Plants, representing gene families that appeared in the ancestors of green algae) to youngest (i.e. *Arabidopsis thaliana*, representing gene families specific to *Arabidopsis thaliana)* based on specific time (termed phylostrata) at which they appeared. We found that the percentage of genes with experimentally verified annotation decreases as the phylostrata become younger (Figure 3C).

To examine if prediction coverage also depends on the gene age, we calculated the percentage of genes with predicted function for each phylostrata. Interestingly, the percentage of genes with predicted function decreases for younger phylostrata (Figure 3C), suggesting that function of younger genes is more difficult to predict. Since Neighbor Counting uses network neighbors with experimentally verified annotation to predict gene function (Figure 1A), we investigated the connectivity of the genes in the different phylostrata. The analysis revealed that younger genes tend to be less connected than older genes, as exemplified for GeneMANIA (Figure 3D) and the remaining networks (Figure S4), explaining why fewer predictions can be made for younger genes (Figure 3C).

### Implementation of an online tool to access the ensemble predictions

To provide easy access to NCE predictions with the aim to facilitate hypothesis generation, we have implemented an online tool, EnsembleNet, available at http://aranet.mpimp-golm.mpg.de/ensemblenet.html. The tool provides three approaches to retrieve functional information of genes: (i) by using a query gene to retrieve other genes with similar function (gene-centric search), (ii) by using a GO term to retrieve genes assigned to the term (gene ontology-centric search), and (iii) by using a group of query genes to retrieve genes with similar function (gene set-centric search). However, to enable these analyses, we had to systematically (i) predict which functions a given gene has and (ii) estimate which genes are functionally related.

To predict gene function, we set to investigate which genes are associated to a given Gene Ontology query term (see Methods). Briefly, (i) NC was used to count the number of networks predicting that given gene is associated to the query term, followed by (ii) a permutation analysis to derive an empirical p-value reflecting how well the prediction could estimate correct GO term-gene associations (exemplified in Figure S5, see Methods). The predicted gene-GO associations are available as Supporting Information File 1. To identify functionally related genes, we used Jaccard index (also called intersection over union) to calculate the similarity of GO terms of any two genes. The Gene Ontology Jaccard Index (GOJI) value ranges from 0 (two genes have no GO terms in common) to 1 (two genes have identical GO terms). We then estimated GOJI>0.178 as a threshold capturing biologically relevant values, as <1% of randomly selected gene pairs are found above this threshold (Figure S6, see Methods). The resulting GOJI network is available as Supporting Information File 2.

The gene-centric search of EnsembleNet starts by entering the ID of a query gene (Figure 4A). The analysis returns a list of genes sorted according to the GOJI value, which indicates how functionally similar the genes are to the query gene. This information is also shown as an interactive GOJI network, which visualizes GOJI relationships between the query gene (large node) and the functionally related genes. Furthermore, a table with the predicted GO terms of the query gene indicates how much evidence (i.e. experimental annotation and co-function network predictions) support this term prediction. The reliability of the predicted GO term-gene association is also indicated by the empirical p-value (exemplified in Figure S5, see Methods). Taken together, the gene-centric analysis can (i) reveal genes that are functionally related to the query and (ii) predict the function of the query.

**Figure 4.**
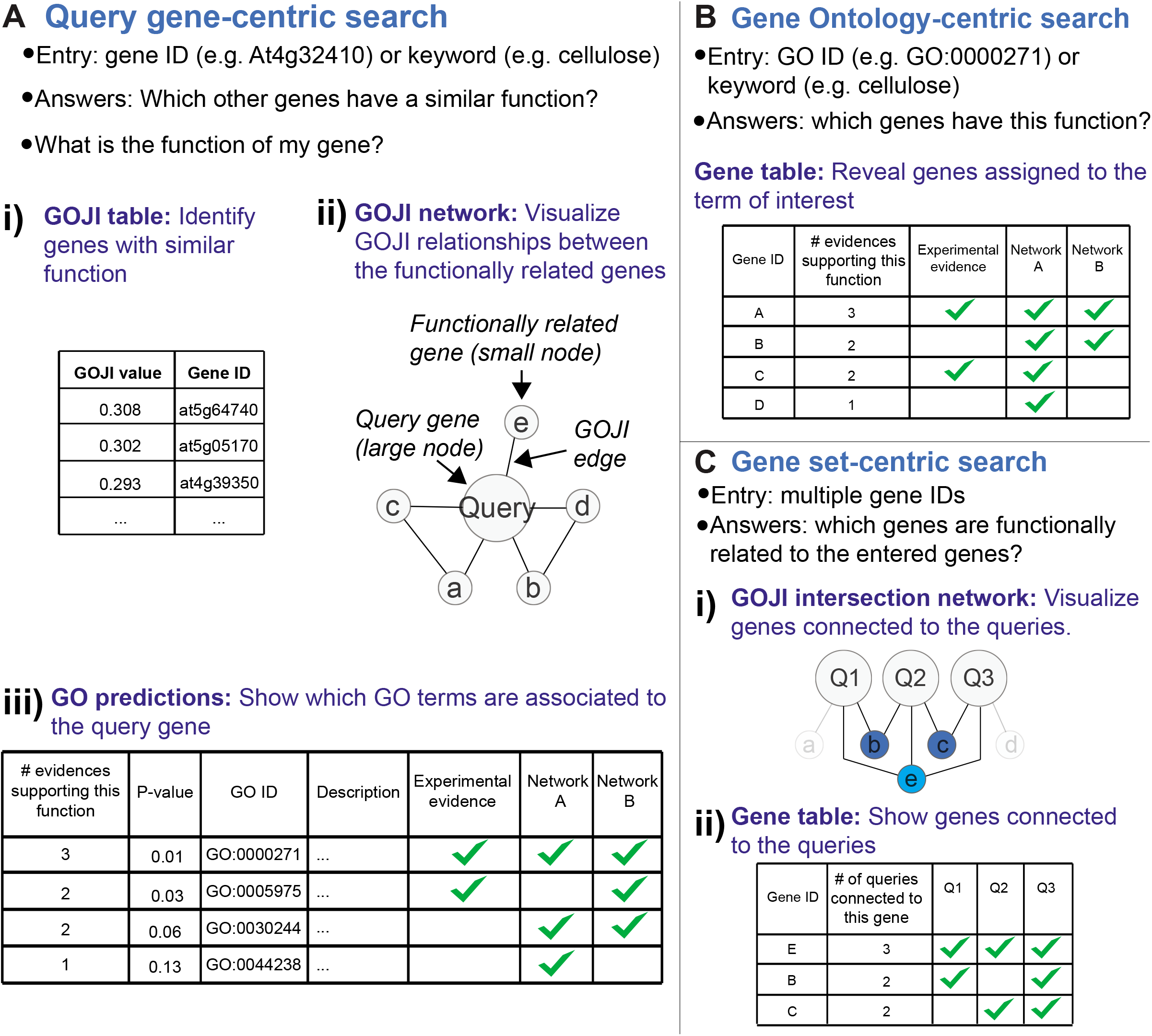
Features of EnsembleNet. A) The query gene-centric can reveal genes that are functionally related to the query, and predict function of the query. Functionally related genes are identified by i) Gene Ontology Jaccard Index (GOJI) table and ii) network. Predicted Gene Ontology (GO) terms of the query are given in the tables below the network. B) Gene Ontology-centric analysis reveals which genes are assigned to a given GO term. For each gene, the table shows how many evidences (experimental annotation and co-function network predictions) support this association. C) Gene set-centric analysis shows i) GOJI intersection network, which reveals genes that are connected to at least two of the query genes. In this example, large nodes correspond to query genes, small nodes represent genes connected to the queries, while edges represent GOJI associations above the significance threshold. Note that genes a and d are removed from the network, as they are connected to only one query gene. ii) Gene table shows to how many queries the found genes are connected to.

The GO-centric analysis reveals which genes are assigned to the query GO term. EnsembleNet first shows the p-value which indicates how well the predictions could correctly predict the genes experimentally associated to the query term, thus indicating the reliability of the predictions for the term. Furthermore, for each gene associated to the term, the table shows how many evidences (experimental and prediction) support this association (Figure 4B). Taken together, the GO-centric search can elucidate novel genes that are involved in a given biological process.

Finally, the gene set-centric analysis accepts several query genes to identify other genes that are functionally related to the queries. The analysis returns a GOJI intersection network, which shows genes that are connected to at least two queries (Figure 4C). By color coding genes according to the number of queries they are connected to, the network can highlights relevant genes that are highly connected to the queries. This information is also presented as a gene table, which is sorted by the number of queries to which a given gene is connected. To summarize, the gene set-centric search can reveal genes that are functionally related to multiple input genes.

### Identification of five complex I assembly mutants

The NADH dehydrogenase complex (complex I) is essential for cellular respiration because it serves as the main site for electron insertion into the mitochondrial electron transport chain, and can provide up to 40% of the protons for mitochondrial ATP formation (Hunte *et al*, 2010; Hirst, 2013; Braun *et al.*, 2014). The number of complex I subunits varies between different eukaryotes due to appearance of some lineage-specific accessory subunits (e.g. *Chlamydomonas reinhardtii:* 42 subunits; *Yarrowia lipolytica:* 42; *Bos taurus:* 45; *Arabidopsis thaliana:* 49) (Cardol *et al.*, 2004; Carroll *et al.*, 2006; Angerer *et al.*, 2011; Peters *et al.*, 2013). Despite decades of research about complex I, new subunit compositions of the complex are still being discovered (Peters *et al.*, 2013; Bridges *et al.*, 2017; Guerrero-Castillo *et al.*, 2017).

We demonstrate how we used the gene-, gene ontology- and gene set-centric approaches of EnsembleNet to identify genes involved in mitochondrial respiratory chain function, and how we arrived at five candidates: *At3g07480, At2g44620, At1g05205, At2g20820* and *At5g64350.* However, any of the three analyses was sufficient to uncover these candidates, and can serve as a stand-alone approach to uncover new, relevant genes.

We first performed a gene-centric search using a NADH dehydrogenase (ubiquinone) gene *At2g02510* (www.gene2function.de/responder.py?name=ens!at2g02510), which is annotated as a subunit of mitochondrial complex I (Klodmann & Braun, 2011). The analysis revealed 174 functionally related genes with GOJI values above the significance threshold (Table S4). While many of these genes are known components of complex I (e.g. *At5g47890, At3g18410, At3g03070*, and others (Klodmann & Braun, 2011), the table also contains several uncharacterized genes, including the five candidates (highlighted in Table S4A). Furthermore, the GOJI network of the top 50 genes contained the five candidates and revealed that most of the genes are fully interconnected, further indicating that these genes are functionally related (Figure 5A, light blue nodes). Finally, the tables below the GOJI network can show predicted BP, CC and MF terms of the query gene. Indeed, respiratory chain complex I (GO:0045271) is one of the top terms found in the CC table of *At2g02510*, supported by all three co-function networks (AraNet v2, ATTED-II and GeneMANIA), and a p-value<0.001 (Table S4B), which is in line with the complex I function of *At2g02510.*

**Figure 5.**
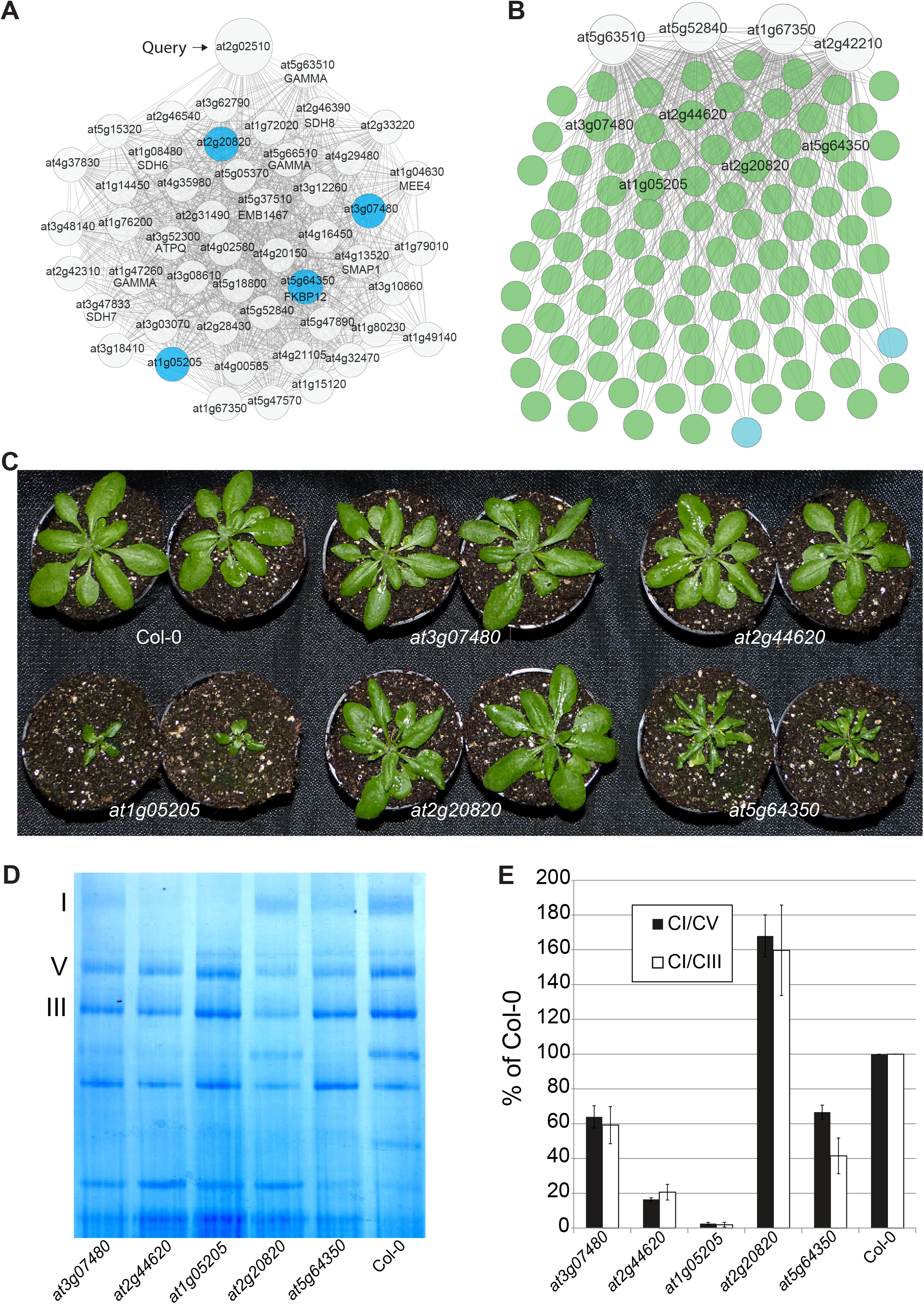
Functional characterization of *at3g07480, at2g44620, atlg05205, at2g20820* and *at5g64350.* A) GOJI network of At4g30010 (large node). Smaller nodes represent genes connected to the At4g30010. The edges connecting the nodes represent GOJI values above the significance threshold. Genes selected for functional characterization are indicated as dark blue nodes. B) Gene set analysis using queries *At5g63510, At5g52840, Atlg67350* and *At2g42210* (large nodes). 100 Genes connected to four (green nodes) or three (blue nodes) of the queries are shown. The five genes selected for functional analysis are indicated. C) Top view of 5 week old rosettes. D) Blue Native Polyacrylamide Gel Electrophoresis (BN-PAGE) analysis of the five mutants and wild type (Col-0). Bands corresponding to complexes I, III and V are indicated to the left. E) Calculated ratio of complex I/complex V (CI/CV) and complex I/complex III (CI/CIII). The bars represent means, while the error bars represent standard deviation (n=4).

Next, we used “mitochondrial respiratory chain complex I” as a term-centric search (www.gene2function.de/responder.py?name=ens!GO:0005747). The analysis returned 1,761 gene candidates, sorted according to the number of evidences (i.e. experimental annotation, and predictions from AraNet v2, ATTED-II and GeneMANIA) supporting the association of a gene to the GO term, which can be used to assign confidence to the prediction. Of the 1761 predictions, 32, 113, 344 and 1272 of the genes were supported by 4, 3, 2 and 1 evidences (Table S5). The five gene candidates were found to be associated to the query term by 2, 3, 3, 2 and 3 evidences for *At3g07480, At2g44620, At1g05205, At2g20820* and *At5g64350*, respectively (Table S5, candidates highlighted in green).

Finally, we conducted a gene set-centric search, by using four genes with experimentally verified association to “mitochondrial respiratory chain complex I”: *At5g63510, At5g52840, At1g67350* and *At2g42210* (Klodmann & Braun, 2011). The analysis revealed genes ranked according to the number of queries they are connected to. For example, the GOJI intersection network of top 100 connected genes showed candidates connected to four (green nodes) or three (blue nodes) queries (Figure 5B, queries shown as large nodes). A table below the network revealed that 98, 108 and 121 genes are connected to 4, 3 and 2 of the queries, respectively (Table S6). The five candidates were connected to the four queries in the GOJI intersection network (Figure 5B, nodes with gene IDs) and the table (Table S6, highlighted genes).

To confirm that the five candidates are involved in complex I formation, we analyzed knock-out mutants of these genes. Phenotypic characterization revealed that two of the mutants showed decreased plant size *(at1g05205* and *at5g64350*, Figure 5C). To see if complex I amount is perturbed in the mutants, we performed a blue native polyacrylamide gel electrophoresis (BN-PAGE) analysis on the mitochondrial proteome (Figure 5D), which allowed us to estimate the abundance of complex I, by calculating the complex I:complex III and complex I:complex V ratios (Figure 5E, Peters *et al.*, 2012). The analysis revealed that the five mutants have significantly different CI/CV and CI/CIII ratios from wild type (Col-0, p-value<0.01, Table S7), where four of the mutants have lower ratio of complex I, while *at2g20820* has higher amount (Figure 5E). Interestingly, while the dwarf phenotypes of *at1g05205* and *at5g64350* are associated with a decrease of complex I ratio, the strong decrease of the ratio in *at2g44620* did not produce any observable growth phenotype (Figure 5C). This was also observed for the *ca2* mutant, where 90% decrease of complex I produced no growth phenotype (Perales *et al.*, 2005).

## Discussion

Gene function prediction is one of the most active fields of bioinformatics, and recent publications show a constant improvement of the predictive power of these approaches (Radivojac *et al.*, 2013; Jiang *et al.*, 2016). In this paper, we studied ensemble gene function prediction, a powerful, but unexplored approach in the plant field.

We used Neighbor counting (Sharan *et al.*, 2007) and the F-measure (Radivojac *et al.*, 2013; Jiang *et al.*, 2016) to benchmark the 10 networks. As expected, the second generation integrative approaches perform better than the first generation networks (Figure 1). However, while the two top networks (GeneMANIA and AraNet v2) use different data sources and methods to integrate them (regression vs. Bayesian, respectively), the performance of the two networks was similar (Figure 1C), suggesting that the performance of the second generation methods might have reached it’s peak.

We next established an ensemble prediction by using two different approaches (Figure 2). The neighbor counting ensemble (NCE) integrates the predictions to arrive at an ensemble prediction, while the STATISnet integrates the co-function networks to arrive at an ensemble network. While integration of the networks via STATISnet produced an inferior prediction than GeneMANIA, NCE showed the highest accuracy and coverage (Figure 2C). In contrasts to conclusions from previous study (Marbach *et al.*, 2012), integration of all 10 methods did not result in highest accuracy, but rather highest prediction coverage at the cost of accuracy (Figure 2C-D). To arrive at the highest accuracy, a combination of GeneMania, AraNet v2 and ATTED-II was needed to address BP and CC, while combination of GeneMANIA and AraNet v2 was better suited to address MF domain. This suggests that careful combination of the inference methods might be needed to achieve highest accuracy for different domains of gene function.

We next investigated how the amount of available information influences the accuracy and coverage of the predictions. Upon removing genes from the gold standard, we observed that the prediction accuracy dropped substantially for gold standard size of <50%, followed by a more modest increase for gold standard size of >50% (Figure 3A, Figure S3). The number of made predictions shows a strong dependency on the size of the gold standard, as in contrast to accuracy, the prediction coverage shows a substantial increase for gold standard size >50%. These results are expected, as function of the uncharacterized genes in this study is transferred from their characterized direct network neighbors, which strongly depends on the fraction of gold standard genes in the network. To partially circumvent this dependency, density-based methods which identify highly connected network modules could be used to annotate genes that are not directly connected (Sharan *et al.*, 2007).

Are functions of certain genes more difficult to predict? Our previous study revealed that older gene families are more functionally characterized than younger gene families (Figure 3C, Ruprecht *et al.*, 2017). Surprisingly, we observed that functions of older gene families can be more readily predicted than for younger genes (Figure 3C), which could be attributed to higher disconnectedness of younger genes (Figure 3D, Figure S4). Interestingly, we observed this in BIOGRID protein-protein interaction networks and to a weaker degree for co-expression networks based on RNA-sequencing data (Figure S4). Higher connectivity of older genes in protein-protein interaction networks could be explained by selection bias (e.g. older genes have stronger mutant phenotypes, and are more likely to be used as bait in yeast-two-hybrid experiments), but we observed the same effect for co-expression networks based on unbiased genome-wide RNA-seq analysis, which indicates that the disconnectedness is an inherent property of young genes. Since it is not possible to predict gene function of disconnected genes, function prediction of the young genes still remains a challenge.

To provide an easy access to the predictions, we present the EnsembleNet database which allows gene-, gene ontology- and gene set-centric searches (Figure 4). These versatile features enabled us to identify 5 uncharacterized genes important for complex I assembly (Figure 5). The high granularity of Gene Ontology terms which allows interrogating the polygenic composition of specific processes (e.g. GO:0035619: “root hair tip”), together with cutting-edge performance, represents a major step towards the goal of computationally bridging the gap between genotype and phenotypes in plants.

## Acknowledgements

We would like to express our gratitude to the creators of ATTED-II, GeneMANIA, STRING, AraNet v2, BIOGRID, PODC and TAIR for providing these excellent tools and networks. We would like to thank the employees of the Max Planck Institute of Molecular Plant Physiology, and especially Dr. Arun Sampathkumar, for their feedback regarding EnsembleNet tool. Marek Mutwil would like to thank Sue Rhee for the great discussions regarding gene prediction function. This work was supported by the Max-Planck-Gesellschaft (B.H, E.M., C.F., N.V., Z.N., M.M.).

## Author contributions

M.M. conceived the project. M.M. and B.H. performed the bioinformatical analyses with input from Z.N.. K.V and S.M. provided CC-AFM network. E.M. performed the mutant analyses. M.M., B.H., E.M., C.F., N.V., K.V. and Z.N. wrote the article with help from all authors.

**Supplemental figures:**

**Figure S1. Status of the functional annotation of the 28392 *Arabidopsis thaliana* genes on November 2015.** A) The pie charts show the fraction of experimentally validated (green), predicted (blue) and unknown function (gray) genes for BP, CC and MF. B) Venn diagram showing the overlap of functionally characterized genes in the three domains of GO.

**Figure S2. Combinatorial analysis of the ten gene co-function networks for BP (top), CC (middle) and MF (bottom).** The x-axis indicates the number of made predictions, while the y-axis shows the F-measure. Point color indicates the number of integrated networks, ranging from 1 network (dark blue) to ten (bright yellow).

**Figure S3. Influence of available functional information on F-measure and the number of made predictions.** Relationship between the F-measure (y-axis) and % of genes with available GO terms (x-axis) given for BP (top left), CC (middle left) and MF (bottom left). Relationship between the number of made predictions (y-axis) and % of genes with available GO terms (x-axis) for BP (top right), CC (middle eight) and MF (bottom right). The lines represent bootstrapped 68% confidence interval plots of 100 permutations.

**Figure S4 Relationship between the percentage of genes with a neighbor in a network (y-axis) and the phylostrata of the genes (x-axis).** The example is given for ATTED-II (top), AraNet v2 (middle) and GeneMANIA (bottom), BP.

**Figure S5. Estimating the performance of Gene Ontology term predictions.** A) An example of a GO term that can be predicted well. Three genes (1-3) experimentally assigned to the term are predicted to be assigned to this term in 9 cases (indicated by green +). B) An example of a GO term that cannot be predicted well. Only 2 out of 9 networks predicted that the experimentally assigned genes A-C are assigned to this term.

**Figure S6 Estimating the significant Gene Ontology Jaccard Index (GOJI) value.** The histogram shows the relationship between GOJI values (x-axis) and frequency of random gene pairs showing the particular value. The red line indicates the GOJI threshold (0.178) below which 99% of the random pairs are found.

**Supplemental tables:**

**Table S1. F-measure, precision, recall, number of made predictions of the 10 cofunction networks at the different edge cutoffs.** The values are given for the three domains of GO. STATISnet is included.

**Table S2. F-measure, precision, recall, number of made predictions for all combinations of the 10 co-function networks at the F_max_ edge cutoffs.** Tables S2A, S2B and S2C correspond to BP, CC and MF, respectively.

**Table S3. Timestamp F-measure, precision, recall, number of made predictions of the 10 co-function networks and NCE.** The method with highest F-measure in a given domain is in bold.

**Table S4. Gene-centric analysis using *At2g02510.*** A) Output of the GOJI table, where genes are sorted according to the decreasing GOJI value. The five candidates are highlighted in green. B) Predicted BP domain terms of *At2g02510.* The number of evidences supporting this function is a sum of evidences (Experimentally verified function, AraNet v2, ATTED-II and GeneMANIA) that support (Y) this association.

**Table S5. GO-centric analysis using *GO:0005747* (“mitochondrial respiratory chain complex I”).** The table is sorted according to the number of evidences (Experimentally verified function, AraNet v2, ATTED-II and GeneMANIA) supporting this association (Y). The five candidates are highlighted in green.

**Table S6. Gene set-centric analysis using queries *At5g63510, At5g52840, At1g67350 and At2g42210.*** The table is sorted according to the number of the queries a given gene is connected to with a GOJI edge. The five candidates are highlighted in green.

**Table S7. Statistical analysis of complex I/V and complex I/III ratios.** The ratios of the mutant and wild type (Col-0) indicate the t-test statistic (n=4).

**Table S8. Primers used to test for wild type allele (RP+LP) and mutant allele (RP+LB) for the five knock out lines.**

**Supporting Information Files:**

**Supporting Information File 1. NCE predicted gene-GO term associations for BP, CC and MF.** The columns in each file show the GO ID, empirical p-value reflecting how well the prediction could estimate correct GO term-gene associations, gene ID, and indication if a given co-function network predicted the association (1), did not predict this association (0), or could not make a prediction for this gene (-).

**Supporting Information File 2. Gene Ontology Jaccard Index (GOJI) network.** The first two columns contain gene IDs, while the third column shows the GOJI value. Only gene pairs above the significance threshold are shown.

